# Cataloging the Phylogenetic Diversity of Human Bladder Bacterial Isolates

**DOI:** 10.1101/2023.05.23.541916

**Authors:** Jingjie Du, Mark Khemmani, Thomas Halverson, Adriana Ene, Roberto Limeira, Lana Tinawi, Baylie R. Hochstedler-Kramer, Melline Fontes Noronha, Catherine Putonti, Alan J. Wolfe

## Abstract

Although the human bladder is reported to harbor unique microbiota, our understanding of how these microbial communities interact with their human hosts is limited, mostly owing to the lack of isolates to test mechanistic hypotheses. Niche-specific bacterial collections and associated reference genome databases have been instrumental in expanding knowledge of the microbiota of other anatomical sites, e.g., the gut and oral cavity. To facilitate genomic, functional, and experimental analyses of the human bladder microbiota, here we present a bladder-specific bacterial reference collection comprised of 1134 genomes. These genomes were culled from bacterial isolates obtained by a metaculturomic method from bladder urine collected by transurethral catheterization. This bladder-specific bacterial reference collection includes 196 different species, including representatives of major aerobes and facultative anaerobes, as well as some anaerobes. It captures 72.2 % of the genera found when we reexamined previously published 16S rRNA gene sequencing of 392 adult female bladder urine samples. Comparative genomic analysis found that the taxonomies and functions of the bladder microbiota shared more similarities with the vaginal microbiota than the gut microbiota. Whole-genome phylogenetic and functional analyses of 186 bladder *E. coli* isolates and 387 gut *E. coli* isolates supports the hypothesis that phylogroup distribution and functions of *E. coli* strains differ dramatically between these two very different niches. This bladder-specific bacterial reference collection is a unique resource that will enable hypothesis-driven bladder microbiota research and comparison to isolates from other anatomical sites.

## Introduction

High-throughput DNA sequencing and enhanced culture-based investigations have found bacterial DNA and live bacteria, respectively, in catheterized (bladder) urine deemed culture-negative by the standard urine culture method^1–13^. The expanded quantitative urine culture (EQUC) protocol and similar enhanced (metaculturomic) methods have enabled researchers to isolate species detected by 16S rRNA gene sequencing in both individuals with and without urinary tract symptoms^7, 8, 11, 13–15^. While the genetic diversity of some species has been extensively investigated, e.g., *Escherichia coli*^16, 17^, many species have few or no genomic sequences. Furthermore, new urinary species are still being discovered, e.g., *Lactobacillus mulieris*^18^, while others are being reclassified in light of new genomic data, e.g., *Gardnerella vaginalis*^19^.

Previously, we published analysis of 149 genomes of 78 different species isolated from the female bladder^14^. Several of the taxa found within the bladder microbiota are also inhabitants of the vaginal microbiota^14, 20, 21^. This led us to posit that the bacterial communities of these two anatomical sites may be connected^14^. A recent 16S rRNA gene sequencing survey of paired bladder and vaginal samples identified more bacterial genera within the bladder than the vagina, suggesting a greater bacterial diversity within the urinary tract^21^. In contrast, little overlap in species was reported between the microbiota of the urinary and gastrointestinal tracts^14^, despite prior evidence that the gastrointestinal tract is the source of *E. coli* that cause urinary tract infections (UTIs)^22–25^. Furthermore, our prior work found that key metabolic pathways specific to the female urogenital environment are not found in bacterial genomes isolated from the gastrointestinal tract, suggesting that these two microbiotas are not connected^14^.

Capturing the species present and the strain diversity of those species is fundamental for the success of gene marker surveys, as well as shotgun metagenomic studies. In contrast to the gastrointestinal tract and oral cavity, very few shotgun metagenomic sequencing studies of the urinary microbiota have been conducted, and then only relatively recently^26–30^. The available genomes representative of the genetic and species diversity within the urinary tract is essential for understanding the urinary microbiota and its association with symptoms and response to treatment. In this study, we combined EQUC with large-scale whole-genome sequencing to generate a comprehensive bladder-specific bacterial genome reference collection. To assess the completeness of this catalog, we compared the taxa found here to those identified in a reexamination of previously published high-throughput 16S rRNA gene sequencing surveys of bladder urine. We compared the genomes of this new culture collection to previously sequenced gut and vaginal isolates and identified taxonomies and functional similarities between the bladder and vagina but not the gut. Finally, we identified phylogenetic and functional differences between *E. coli* strains isolated from the bladder and gut.

## Results

### Demographics

In an effort to assemble a genomic reference collection representative of the phylogenetic diversity of the bladder, participants of this study were recruited as part of previous and ongoing IRB-approved studies. Out of a total of 5619 bladder isolates in our urinary bacteria collection, we selected 1050 representatives for whole genome sequencing. Some isolates came from the same participant. In those cases, the isolates were always from different species.

These representative bladder strains were isolated, using a metaculturomic method called Expanded Quantitative Urine Culture (EQUC)^7, 8^, from urine obtained by transurethral catheterization of 96 asymptomatic controls and 377 symptomatic individuals. The symptomatic individuals had been diagnosed with urinary tract infection (UTI, n=212), recurrent urinary tract infection (RUTI, n=54), overactive bladder (OAB, n=76), stress urinary incontinence (SUI, n=10), bladder cancer (n=12), bladder and bowel dysfunction (BBD, n=3), kidney stone (n=1), interstitial cystitis/bladder pain syndrome (IC/BPS, n=1), pelvic organ prolapse (POP, n=5) or unknown urinary tract symptoms (n=3). (n=3). Participants were from 6 self-reported races: White/Caucasian (n=340), Black (n=57), Hispanic (n=41), Asian (n=5), Native American (n=3), Middle Eastern (n=2), and no reply (n=25). Most participants were females (n=454); only 19 were males. Participants ranged in age from 4 weeks to 99 years old (**Table 1**). Participant metadata for each genome are listed in **Supplementary Table 1**.

**Table 1:** Demographics of participants in this study.

|  |  | Symptomatic individuals<br>(n=377) | Asymptomatic controls<br>(n=96) |
| --- | --- | --- | --- |
| **Sex** | Female | 366 | 88 |
|  | Male | 11 | 8 |
| **Age** | <18 | 2 | 5 |
|  | 18-40 | 28 | 23 |
|  | 40-65 | 109 | 45 |
|  | >65 | 227 | 15 |
| **Disease states** | UTI | 212 | N/A |
|  | RUTI | 54 | N/A |
|  | OAB | 76 | N/A |
|  | SUI | 10 | N/A |
|  | Bladder cancer | 12 | N/A |
|  | BBD | 3 | N/A |
|  | Kidney stone | 1 | N/A |
|  | IC/BPS | 1 | N/A |
|  | POP | 5 | N/A |
|  | With symptoms | 3 | N/A |
| **Self-reported Race** | White | 282 | 58 |
|  | Black | 41 | 16 |
|  | Hispanic | 29 | 12 |
|  | Asian | 5 | 0 |
|  | Native American | 2 | 1 |
|  | Middle Eastern | 2 | 0 |

### The bladder-specific isolate genomes are diverse and large

Despite best efforts, many clinical isolates cannot be completely purified. Besides the primary, originally identified species, some isolates contained one or more secondary, minority species; in another system, these minority species have been called hitchhikers^31^. Of our 1050 isolates, 218 consisted of more than one species. In total, we obtained 1134 high-quality genomes (>90% completeness, <5% contamination), including 781 genomes produced as part of this current effort and 353 genomes from our collection that were previously deposited (**Supplementary Figure 1, Supplementary Table 1**). The overall qualities of these genome assemblies were high: the median completeness level was 99.39% and the median estimated contamination was 0.86%. The median positional coverage was 160× and the median contigs L50 (the number of contigs needed to cover 50% of the genome assembly) was 14.24 (**Fig. 1A–D**).

**Figure 1:**
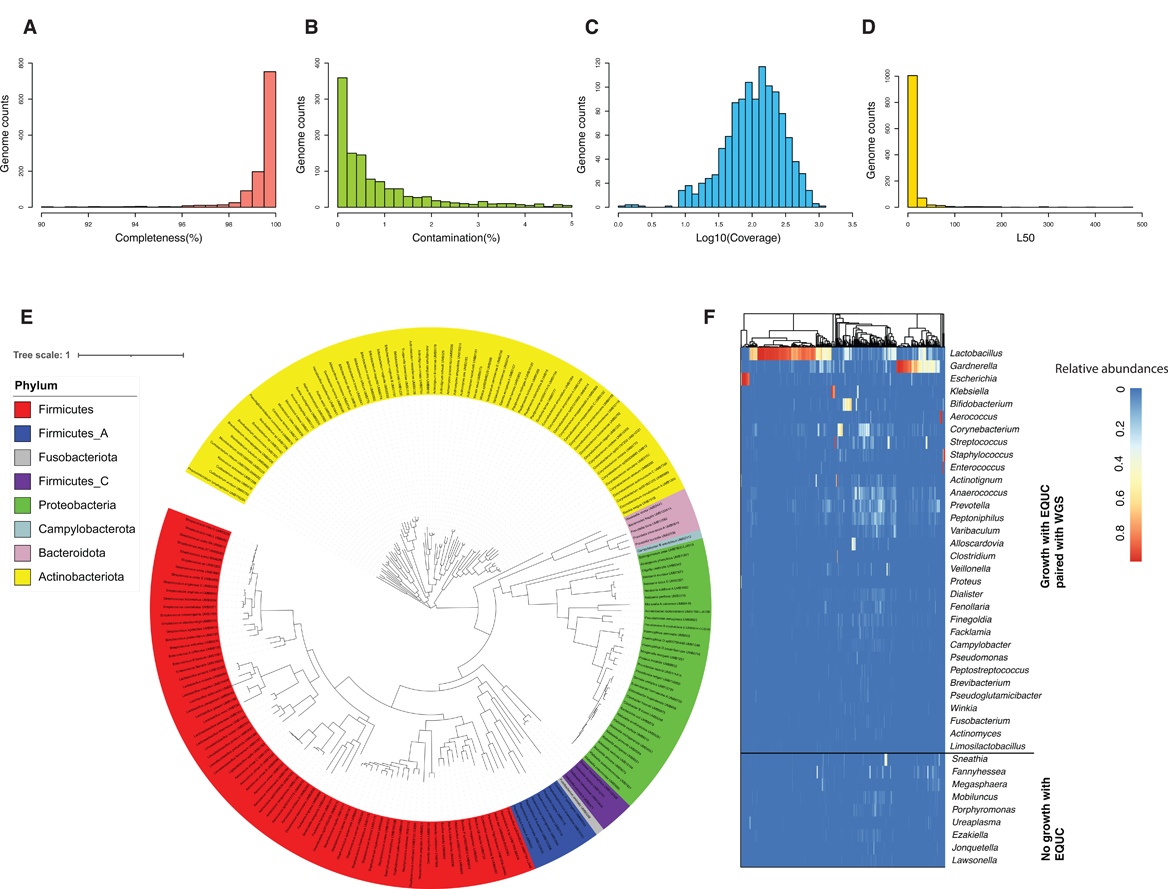
Overall information of the bladder microbiota culture collection. A-D. Distribution of genome assembly, completeness, contamination, coverage (on a log10 scale), and L50 values, respectively. E. Phylogenomic tree of the 196 bacterial species (assigned by GTDB) represented in the bladder genome collection. A single representative genome per species was selected out of the 1134 high quality isolated genomes (>90% completeness, < 5% contamination) to reconstruct the multiple sequence alignment of 71 core bacterial genes. Bacterial species are colored by phylum. F. The diversity of isolate genomes was compared with the total diversity observed in the 16S sequence data (as shown in genus level) obtained from 392 bladder urine samples.

Most genomes (n=1096) were from females; only 38 were from males (**Supplementary Table 1**). This collection of genomes represented 8 phyla, 10 classes, 21 orders, 39 families, 89 genera, and 196 species (**Figure 1E**). This includes 7 genomes for which neither the NCBI taxonomy database nor the Genome Taxonomy Database (GTDB) could assign a species (**Supplemental Table 1**). These 7 genomes (from UMB1203, UMB1308A, UMB1298, UMB7805-LC452B, UMB4589-SE434, UMB1308B, and UMB10442) represent isolates of the genera *Brevibacterium*, *Corynebacterium, Paenibacillus_B*., *Sphingomonas*, and *Streptococcus*.

Pairwise whole genome average nucleotide identity (ANI) comparison of the 196 species met a 95%-96% threshold (**Supplementary Figure 2)**, the generally accepted ANI threshold for distinguishing between two bacterial species (94%-96%)^32, 33^. More species were isolated from symptomatic participants (n=169, **Supplementary Figure 3**) than from asymptomatic participants (n=76, **Supplementary Figure 4**). This skew likely resulted for two reasons: (1) 79% of the genomes were from symptomatic participants and (2) overall, symptomatic participants tend to have richer urinary bacterial communities (i.e., possess more unique species)^34^.

To determine how well our genome collection represents the bacterial diversity captured by culture-independent methods, we re-analyzed 16S rRNA gene sequencing data previously obtained from 392 female bladder urine samples^3, 5, 35–37^ (**Supplemental Table 2**). We found that the sequenced genomes represented 72.2% (26/36) of the most abundant genera (>0.1% relative abundance) captured by 16S sequence analysis (**Fig. 1F**). Nine genera were detected by 16S rRNA gene sequencing without representative strains within our genome collection: *Sneathia*, *Fannyhessea*, *Magasphaera*, *Mobiluncus*, *Porphyromonas*, *Ureaplasma*, *Ezakiella*, *Jonquetella*, and *Lawsonella*. These genera are strictly anaerobic, particularly fastidious, or lack a cell wall.

For other abundant anaerobes detected by 16S sequencing, such as *Anaerococcus* and *Prevotella*, we had only a few strains in our cultured isolate collection (**Supplementary Figure 1, Supplementary Table 1**). For example, the average relative abundance of *Anaerococcus* and *Prevotella* is about 2.8% and 3.2%, respectively, among all 16S rRNA gene sequenced samples, but our collection of 5619 isolates only contains 3 *Anaerococcus* and 5 *Prevotella* strains.

In contrast, we were able to isolate and thus sequence taxa that were simply missed by 16S rRNA gene sequencing or were present only as rare taxa (<0.1% average relative abundances across all individuals); these included isolates from the genera *Acinetobacter*, *Citrobacter*, *Cutibacterium*, *Enterobacter*, *Gleimia*, *Haemophilus*, *Micrococcus*, *Neisseria*, and *Oligella* (**Supplementary Tables 1 and 2**).

### Bladder isolates are functionally and taxonomically more similar to vaginal isolates than gut isolates

To assess the bladder microbiota relative to those of other well-studied anatomical sites, we compared the bladder strains we had isolated from asymptomatic females (n=69) to gastrointestinal and vaginal strains isolated by others from unrelated asymptomatic individuals. We computed the ANI between representative genomes of the 69 bladder bacterial species isolated from females to representative genomes of 74 vaginal bacterial species (including 4 species from our vaginal isolate collection) and representative genomes of 175 gut bacterial species, all cultivated from unrelated asymptomatic individuals (**Supplementary Table 3**). Only 5 species (*Bifidobacterium bifidum*, *Enterococcus faecalis*, *E. coli*, *Lacticaseibacillus rhamnosus* and *Proteus mirabilis*) were detected in all 3 niches, and only 9 species were found in both the bladder and gut microbiota. In contrast, 19 species were found in both the bladder and vaginal microbiota (**Figure 2A, Supplementary Table 3**). Taken together, these results indicate that bacterial species from each niche are distinct, but the bladder shares more bacterial species with the vagina than the gut.

**Figure 2:**
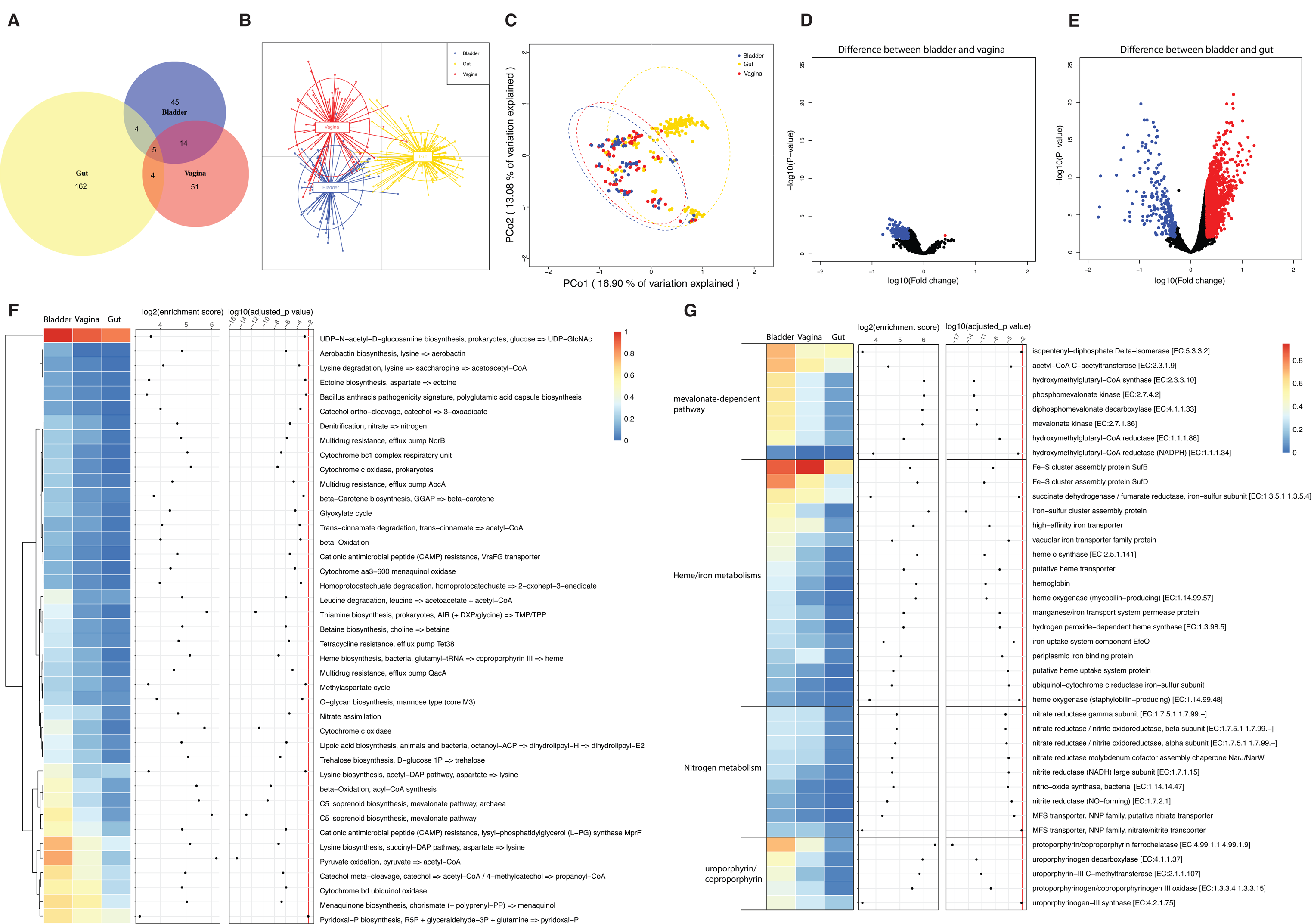
Comparison of bladder isolates with vaginal and gut isolates. A. Venn diagram showing the number of bacterial species isolated from asymptomatic individuals shared among 3 different niches: bladder (red; n=68), vagina (blue; n=74) and gut (green; n=175). Reference genomes of vagina and gut isolated from asymptomatic donors were obtained from previous studies. Bladder isolates from female asymptomatic controls were used in the comparison. B-C. DAPC and PCoA of KEGG functional diversity of representative bacterial species isolated from the asymptomatic individuals of different niches. D-E. Volcano plots of *t* tests corrected by the Benjamini and Hochberg method for changes of KOfam functions between bladder and vagina and between bladder and gut. An FDR cutoff of 0.01 and [logFC]>0.3 were used. Data points highlighted in red indicate KOfam functions that were significantly enriched in gut or vagina, while data points highlighted in blue indicate KOfam functions that were significantly enriched in the bladder. F. KO modules that were enriched in the bladder compared with the gut and/or vagina. An FDR cutoff of 0.01 and more than 10% prevalence in all bladder genomes were used. G. Selected KOfam functions that were enriched in the bladder as compared with gut and/or vagina. Functions that had at least 10% prevalence in all bladder genomes with an FDR cutoff of 0.01 were used.

We next evaluated functional differences in the microbiota by analyzing the protein functions encoded by the representative bacterial species in the three niches. Applying Discriminant Analysis of Principal Components (DAPC) and Principal Coordinates Analysis (PCoA) using the Bray-Curtis Dissimilarity Index, we compared the Kyoto Encyclopedia of Genes and Genomes (KEGG, **Fig. 2B-C, Supplementary Table 4**) and Clusters of Orthologous Groups (COG, **Supplementary Figure 5A-B, Supplementary Table 5**) annotations of the 69 bladder, 74 vaginal, and 175 gut bacterial species. Like the species analysis described above, this comparison identified more overlapping protein functions shared by the bladder and vaginal bacterial species that were largely distinct from protein functions found in the gut bacterial species. Volcano plots also revealed 321 KOfam functions differentially abundant between bladder and vaginal species (t-test, adjusted-p<0.01, [logFD]>0.3, **Figure 2D)**, and 1778 KOfam functions differentially abundant between bladder and gut species (t-test, adjusted-p<0.01, [logFD]>0.3, **Figure 2E**). We observed similar findings for COG functions. 115 COG functions were differentially abundant between bladder and vaginal species. In contrast, 1611 COG functions were differentially abundant between bladder and gut species (t-test, adjusted-p<0.01, [logFD]>0.3, **Supplementary Figure 5C-D**). These data indicate the existence of many functions shared by bladder and vaginal bacterial species that are clearly distinct from those of gut bacterial species.

We identified Kegg Orthology (KO) modules that were significantly different between the 3 niches (**Supplementary Table 6**). Forty-one KO modules were enriched in the bladder (adjusted-p<0.01 and >10% prevalence in all bladder genomes) compared with the gut and/or vagina (**Fig. 2F**). For example, the mevalonate-dependent pathway for isoprenoid biosynthesis was enriched in the bladder genomes (**Fig. 2F, Supplementary Table 6**). All KOfam functions associated with the mevalonate-dependent pathway were enriched in the bladder genomes (**Figure 2G and Supplementary Table 7**), including acetyl-CoA C-acetyltransferase, hydroxymethylglutaryl-CoA synthase, hydroxymethylglutaryl-CoA reductase (NADPH), hydroxymethylglutaryl-CoA reductase, mevalonate kinase, phosphomevalonate kinase, diphosphomevalonate decarboxylase and isopentenyl-diphosphate Delta-isomerase. We also observed enriched KO modules in bladder genomes related to acetyl-CoA, which is the precursor for the MEV pathway of isoprenoid biosynthesis. These acetyl-CoA-related KO modules include beta-oxidation, acyl-CoA synthesis, pyruvate oxidation. Some bladder-enriched KO modules relate to lysine metabolism (i.e., lysine degradation and biosynthesis) or nitrogen metabolism (i.e., denitrification and nitrate assimilation), while others relate to iron utilization (e.g., aerobactin biosynthesis and heme biosynthesis) (**Figure 2F, Supplementary Table 6**). Given the scarcity of iron in the bladder, we investigated specific functions involved in iron utilization/assimilation. Seventeen KOfam functions related to heme biosynthesis and degradation, as well as to iron utilization and transport, were enriched in bladder genomes, including iron−sulfur cluster assembly proteins, hemoglobin, heme o synthase, and iron uptake system component EfeO among others. Several bladder-enriched functions were related to protoporphyrin/coproporphyrin utilization, including protoporphyrin/coproporphyrin ferrochelatase, which uses ferrous iron as one of its substrates (**Figure 2G and Supplementary Table 7**). Analyses of COG functions also supported the enrichment of the mevalonate-dependent pathway, enterochelin transport and pyruvate oxidation in bladder genomes (**Supplementary Figure 5E, Supplementary Figure 6, Supplementary Table 8**). Enrichment of the mevalonate-dependent pathway over the more common methylerthritol 4-phosphate pathway and of functions involved in iron utilization and export may relate to iron availability within the bladder.

### Genomes of *Escherichia coli* strains isolated from the bladder differ from the gut

Given the significant taxonomic and functional differences between bladder and gut genomes in asymptomatic individuals, we next sought to evaluate whether isolates of the same species differed in these 2 niches. As the gut is generally considered to be the source of uropathogenic *Escherichia coli* (UPEC) strains^38, 39^, we compared our 186 bladder *E. coli* genomes with 387 publicly available gut *E. coli* isolates from unrelated healthy individuals^40^.

From the genomes of these 186 bladder isolates, we constructed a phylogenomic tree based on 1084 single-copy core genes (**Figure 3A**). These genomes belonged to two distinct clades and six phylogroups. Phylogroup B2 and one unknown phylogroup belonged to clade 1, whereas phylogroups G, F, A, B1 and D belonged to clade 2. Phylogroup B2 (58.6%, n=109) predominated, followed by phylogroups D (19.4%, n=36), B1 (9.7%, n=18), A (8%, n=15), F (3.2%, n=6), G (0.5%, n=1), and an unknown phylogroup (0.5%, n=1) (**Table 2, Supplementary Table 9**). These genomes were from *E. coli* strains isolated from the bladders of 11 asymptomatic controls and 175 symptomatic individuals who were diagnosed with a urinary tract infection (UTI, n=130), recurrent UTI (RUTI, n=14), overactive bladder (OAB, n=21), stress urinary incontinence (SUI, n=2), bladder cancer (n=4), bladder bowel disorder (BBD, n=2), Interstitial Cystitis/Painful Bladder Syndrome (IC/PBS, n=1) or Pelvic Organ Prolapse (POP, n=1). However, symptom status was not associated with either clade or phylogroup (**Figure 3A**). The gut genomes also belonged to the same 6 phylogroups; however, the distribution differed. In contrast to the bladder isolates, only 3% of the gut genomes belonged to phylogroup B2 (n=13) while 33.85% belonged to phylogroup D (n= 131), 37% to phylogroup F (n=143), 23.5% to A (n=91), 1.8% to B1 (n=7), 0.26% to phylogroup G (n=1), and 0.26% to an unknown phylogroup (n=1) (**Table 2, Supplementary Table 9**). Thus, phylogroup distribution differed dramatically between these 2 niches.

**Figure 3:**
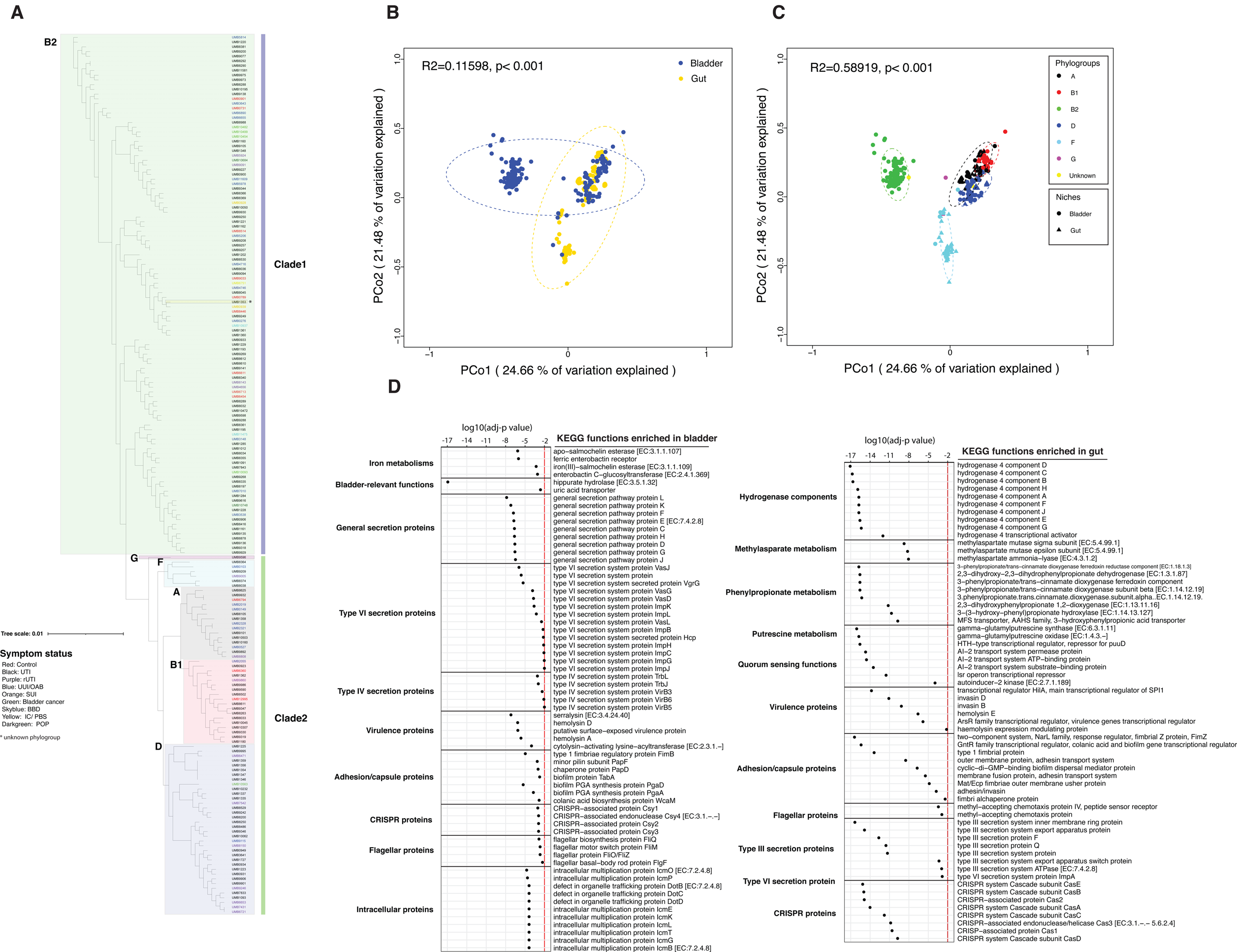
Bladder-specific *Escherichia coli* reference genomes different from the gut reference genomes. A. Phylogenomic relationship of 186 *E. coli* isolates from the bladder derived using 201624 single-copy core genes from their genomes. Phylogroups are indicated by shaded colors. The symptom status of the host for each isolate is indicated by the color of the isolate name. B. Functions of *E. coli* genomes annotated by KEGG are separated by niches. C. Functions of *E. coli* genomes annotated by KEGG are separated by phylogroups. D. Overrepresented KOfam functions associated with bladder-specific or gut-specific genes.

**Table 2:** Number and percent of different *E. coli* phylogroups in bladder and gut genomes.

|  | B2 | A | B1 | D | F | G | Unknown |
| --- | --- | --- | --- | --- | --- | --- | --- |
| Bladder | 109 (58.6%) | 15 (8%) | 18 (9.7%) | 36 (19.4%) | 6 (3.2%) | 1 (0.5%) | 1 (0.5%) |
| Gut | 13 (3.4%) | 91 (23.5%) | 7 (1.8%) | 131(33.9%) | 143 (37%) | 1 (0.26%) | 1 (0.26%) |

We next assessed functions. Applying PCoA using the Bray-Curtis Dissimilarity Index, we compared the annotated KEGG functions of the bladder (**Supplementary Table 10**) and gut *E. coli* genomes (**Supplementary Table 11**) and found that they differed significantly (**Figure 3B**, R^2^= 0.11598, p<0.001). This difference may be because the functions of different phylogroups significantly differed (**Figure 3C**, R^2^=0.58919, p<0.001), especially between phylogroup B2 and the other phylogroups, and that the distribution of the phylogroups in bladder and gut differed (**Table 2**).

We next identified functions that may facilitate *E. coli* adaptation and colonization to these 2 environmentally different niches by evaluating predicted functions encoded by the bladder and gut genomes that were both abundant and differentially present (**Fig. 3D, Supplementary Table 12**). Bladder-enriched functions included those associated with iron metabolism (i.e., the siderophores salmochelin and enterobactin), bladder-relevant functions (i.e., hippuric hydrolase and a uric acid transporter), the general secretion pathway, both type VI and IV secretion systems, and intracellular multiplication and trafficking proteins. In contrast, gut-enriched functions related to hydrogenase components, methylasparate, phenylpropionate metabolism, putrescine metabolism, type III secretion system, and quorum sensing. Both sets of genomes harbored different enriched functions related to virulence, flagella, capsule, adhesion, and CRISPR.

To further evaluate the virulence and antibiotic susceptibility profiles of *E. coli* strains in these 2 niches, we mapped the *E. coli* genomes against the Virulence Factor Database (VFDB) and Comprehensive Antibiotic Resistance Database (CARD). 187 different virulence factors and 53 antibiotic resistance genes were detected in the 186 bladder genomes **(Supplementary Tables 13 and 14)**, and 128 different virulence factors and 16 antibiotic resistance genes were detected in the 387 gut genomes **(Supplementary Tables 15 and 16)**. The number of virulence and antibiotic-resistance genes was significantly greater in the bladder *E. coli* genomes (**Supplementary Figure 7A**). Again, this may result from the distribution of phylogroups, as phylogroup B2 (average counts per genome = 84.3) had slightly more virulence and antibiotic resistance genes than phylogroups D (average counts per genome = 79.6) and F (average counts per genome = 81.7), and much more than phylogroups A (average counts per genome = 67.65) and B1 (average counts per genome = 57.88) **(Supplementary Figure 7B)**.

Seventy-seven virulence and antibiotic resistance genes were enriched in the bladder genomes (**Supplementary Figure 7C, Supplementary Table 17**), whereas 35 were enriched in the gut genomes (Wilcoxon rank sum test, adj-p<0.01, **Supplementary Figure 7D, Supplementary Table 17**). For example, the virulence factor haemoglobin protease (*vat*) and the antibiotic resistance gene cephalosporin-hydrolyzing class C beta-lactamase EC-5 (*blaEC-5*) were present in 44% and 55% of bladder genomes, respectively, but only in 0.5% and 0.3% of gut genomes, respectively. Other virulence genes enriched in the bladder genomes included those that are predicted to encode adhesins (e.g., *afaE-V, focA, C-D, F-H, papA-F, H*, and *sfaB-H, S, X-Y*), heme/iron related virulence (e.g., *chuA, S-T, X-Y, iroB-E, N, irp1-2*, and *ybtA, E, P-Q, S-U, X*), toxins (e.g., *cnf1, hlyA-D, ibeA*, and *tcpC*), and resistance to several antibiotics (e.g., aminoglycosides, beta-lactamases, macrolides, tetracycline, and trimethoprim). In contrast, the gut strains were highly enriched for genes associated with type III secretion and enriched for multiple other virulence factors, including adhesion genes (e.g., *faeC-J, fdeC*, and *ecpC-E*), heme/iron related metabolism (i.e., *iucA-D* and *shuA, T, X*), toxins (i.e., *sat* and *astA*), and 3 class C beta-lactamase genes. Taken together, these results suggest that bladder *E. coli* strains from asymptomatic individuals harbor more virulence and antibiotic resistance genes than commensal gut *E. coli* strains and that *E. coli* strains use different virulence genes to adapt to different niches.

## Discussion

Here, we have presented the largest collection of bladder-specific bacterial isolates paired with whole-genome sequences, providing a valuable resource for hypothesis-driven bladder microbiome research. This extensive, genome-sequenced culture collection represents more than 70% of bacterial genera detected by previously reported 16S rRNA gene sequencing of bladder urine samples collected by transurethral catheterization^3, 5, 35–37^. The missing genera are primarily strict anaerobes that are not aerotolerant, but also include a few that are particularly fastidious or lack a cell wall. For example, per 16S rRNA gene or shotgun metagenomic sequencing, the bacterial genera *Rothia*, *Dialister*, *Bacteroides, Peptoniphilus, Prevotella*, and *Anaerococcus* are abundant and prevalent in catheterized urine samples^4, 20, 35, 41–45^; however, we have few or no isolates from these genera in our collection. Efforts to obtain more representatives of aerotolerant anaerobes should be encouraged. Methods for the culture of non-aerotolerant anaerobes in urine samples also should be developed; this would require a method that does not expose the urine sample to air during collection.

The catalog reported here is a substantial extension of a previously published study^14^. In that previous study, we analyzed 149 bladder genomes from 78 different species. In the current study, we analyzed 1134 genomes from 196 species (**Figure 1E, Supplemental Figure 1**), including 7 species without representative genomes in GTDB or the NCBI taxonomy database. With more comprehensive collections of genomes from the bladder, vagina, and gut, we found the functions and taxonomies of bladder isolates to be more like vaginal isolates than to gut isolates (**Figure 2**), consistent with the previous report^14^. For example, both studies observed enrichment in the bladder of the mevalonate-dependent pathway for isoprenoid biosynthesis over the more common methylerthritol 4-phosphate pathway. This enrichment primarily results from the presence of certain bladder-specific species in the genera *Aerococcus, Actinotignum, Bifidobacterium, Corynebacterium, Enterococcus*, *Lacticaseibacillus, Lactobacillus, Limosilactobacillus, Staphylococcus*, and *Streptococcus.* Other bladder-enriched functions include several related to acetyl-CoA, the precursor to the mevalonate-dependent pathway, and several involved in iron utilization and export. The latter is likely related to the scarcity of iron in the bladder. The reason for enrichment of functions related to lysine metabolism are less obvious.

Since the genomes of all the isolates from bladder and gut differed substantially, we wondered whether the same might be true of a single species found in both niches. We chose *E. coli* for three reasons: (1) it is the most common cause of UTI, (2) it is the most fully characterized bacterial species, and (3) it was the species with the largest number of sequenced genomes from the bladder. Our analysis identified both phylogenetic and functional differences. For example, the bladder and gut genomes belonged to the same 6 phylogroups; however, their distribution differed with phylogroup B2 most common in the bladder, but A, D and F most common in the gut. In fact, phylogroup B2 was relatively rare (3%) in the asymptomatic gut samples examined here. Previous investigation of *E. coli* phylogroup diversity in the gut found the presence of B2 strains most prevalent in individuals with inflammatory bowel disease^46^. Prior studies suggested that most *E. coli* strains causing UTI and other extraintestinal infections belong to phylogroup B2, presumably due to its greater pathogenetic capacity relative to the other phylogroups^47, 48^. However, we did not find an association between symptom status and phylogroup. Although most bladder *E. coli* isolates collected from participants diagnosed with UTI or RUTI belonged to phylogroup B2, this also was true for bladder isolates from asymptomatic controls. These results support the hypothesis that UTI symptoms most likely result from multiple factors, including the composition of the rest of the bladder microbiome and the host response^49–53^.

The difference in phylogroup distribution suggests that only a subset of gut *E. coli* strains are adapted for life in the bladder. From enrichment analysis, it appears that *E. coli* uses different strategies to adapt to and colonize these 2 environmentally different niches (**Figure 3**). Bladder-enriched functions included those associated with iron metabolism, hippuric acid hydrolase, and a uric acid transporter. Given the scarcity of iron and the presence of hippuric and uric acids in the bladder^54^, this makes sense. In contrast, gut-enriched functions related to hydrogenase components, the methylasparate cycle, phenylpropionate metabolism, putrescine metabolism and quorum sensing. These are functions expected of strains adapted for survival in the highly crowded, anaerobic environment of the gut. The observation that the bladder genomes were enriched for type IV and VI secretion systems, as well as intracellular multiplication and trafficking proteins, whereas the gut genomes were enriched for type III secretion systems also suggests very different lifestyles.

### Concluding Remarks

Comparisons between genomes isolated from the bladder, vagina, and gut show that the genetic content of bacteria that inhabit the bladder are distinct, most notably by the functionalities they encode. With the genomes produced through this study, a more comprehensive catalog of bladder bacteria species is now available, representative of >70% of bladder genera detected via high throughput 16S rRNA sequencing surveys. Most notably, the 1134 strains examined here capture the genetic diversity of key bladder species associated with both symptoms and the lack thereof. The genomes, coupled with the participant metadata, provides a key reference for future hypothesis-driven research into the urinary microbiota.

## Methods

### Study design and sample collection

Following Institutional Review Board approval from Loyola University Medical Center (LUMC), University of California San Diego (UCSD) Health, University of Iowa, University of California San Francisco (UCSF) Medical Center, Nationwide Children’s Hospital (NCH) or University of Pittsburgh (Pitt) School of Medicine, participants gave verbal and written consent for chart abstraction and urine collection with analysis for research purposes. Urine samples were collected via transurethral catheter. Urine samples were placed into BD Vacutainer^®^ plus C&S preservative tubes (Becton, Dickinson and Co., Franklin Lakes, NJ). All specimens from LUMC were transported within 4 hours to the Wolfe lab at Loyola University Chicago. Specimens from elsewhere were shipped overnight.

### Expanded quantitative urine culture (EQUC)

EQUC was performed as previously described^7, 8^. Briefly, 100 µl of catharized urine was grown under five conditions with BD BBL prepared plated media: (1) Blood Agar Plate (BAP) in 5% CO_2_ for 48 h, (2) chocolate agar (CHOC) in 5% CO_2_ for 48 h, (3) colistin and nalidixic acid (CNA) agar in 5% CO_2_ for 48 h, (4) CDC anaerobe BAP in an anaerobic jar (BD GasPak Anaerobe Sachets) or anerobic chamber (Coylabs) for 48 h, and (5) BAP under aerobic conditions for 48 h, and MacConkey under aerobic or 5% CO_2_ for 48 h, all at 36°C. In one study (FUN) EQUC was modified to select for fungal microbes as well: BD BBL prepared plated media (1) Brain Heart Infusion Agar, with 10% Sheep Blood, Gentamicin, and Chloramphenicol Gentamicin-BHI in 5% CO_2_, (2) Hardy Chrome-Candida (Chrom-Candida), and (3) Inhibitory Mold Agar (IMA) for 120 h, all at 36°C. The detection level was 10 CFU/ml, represented by 1 colony of growth on any of the plates. In another study (PBSA), EQUC was modified to include BD BBL plate media, Thayer Martin agar for 48 h at 36°C. Each morphologically distinct colony type was isolated on a different plate of the same medium to prepare a pure culture that was used for identification using Matrix-assisted laser desorption/ionization-time of flight (MALDI-TOF) mass spectrometry (MS).

### DNA extraction and whole-genome sequencing

353 genomes from our collection were sequenced and reported previously, as described^14, 55–88^. The remaining 781 genomes were produced as part of this current effort. Their isolates were grown in their preferred medium and pelleted. To extract genomic DNA, cells were resuspended in 0.5 mL DNA extraction buffer (20 mM Tris-Cl, 2 mM EDTA, 1.2% Triton X-100, pH 8) followed by addition of 50 µL lysozyme (20 mg/mL) and 30 µL mutanolysin (5 kU/mL, resulting in 0.15 kU/sample). After a 1 h incubation at 37°C, 80 µL 10% SDS and 20 µL proteinase K were added, followed by a 2 h incubation at 55°C. From this point genomic DNA was either purified with with phenol-chloroform or the MagMax DNA Multi-Sample Kit, according to manufacturer instructions. For phenol-chloroform extractions, 210 µL of 6 M NaCl and 700 µL phenol-chloroform were added. After a 1 h incubation with rotation, the solution was centrifuged at 13,500 rpm for 10 m, and the aqueous phase extracted. An equivalent volume of isopropanol was added; after a 10 m incubation, the solution was centrifuged at 13,500 rpm for 10 m. The supernatant was decanted, and the DNA pellet precipitated using 600 µL 70% ethanol. Following ethanol evaporation, the DNA pellet was resuspended in nuclease-free H_2_O and stored at −20°C. For both approaches, DNA purity and quality were spot checked with Nanodrop Spectrophotometer. 1% agarose gel electrophoresis was performed in difficult-to-extract isolates to confirm genomic DNA isolation and assess degradation. DNA was quantified with the Qubit Fluorimeter Broad Range or with High Sensitivity Kits, depending on yield.

To sequence, the samples were normalized to a maximum of 16ng/uL and library prepped with the Illumina Nextera Flex library prep kit with Nextera XT Indexes. 193 samples were prepared using the Qiagen FX library prep kit with Qiagen Indices. Libraries were quantified with qubit, size distribution assessed with Agilent Bioanalyzer HS kit, and were pooled. A quality control PE150 MiSeq flow cell was run on the pools. Successful pools were sent to the Northwestern University Core Sequencing Facility, where they were sequenced on a Novaseq 6000 yielding PE150 reads for an approximate target of 50x coverage.

### Whole-genome sequence analysis and annotation

The quality of the raw reads was assessed using FastQC (https://www.bioinformatics.babraham.ac.uk/projects/fastqc/). Then, the raw reads were trimmed and filtered using BBMap in BBTools (v38.94) (https://jgi.doe.gov/data-and-tools/bbtools/). Adaptors in samples were first removed using reference (adapters.fa), and trimmed using default settings (ktrim=r, k=23, mink=11, hdist=1, tpe, tbo). Bases with low quality scores (qtrim=rl, trimq=20) and positions with high compositional bias (ftl=20, ftr=135) were removed from both ends, keeping only reads with lengths above 30 bp (minlength=30), with no Ns (maxns=0), and with an average quality above 20 (maq=20). After quality control, all the clean paired-end reads were assembled using SPAdes (v3.14.1)^89^ specifying the (--isolate) mode with full k-mer size list (-k 21,33,55,77). To assess completeness and contamination of the assembled genomes, CheckM (v1.0.12)^90^ was performed using the ‘lineage_wf’ pipeline, and genomes were filtered at completeness ≥90% and contamination ≤5% to obtain high-quality genomes. MaxBin 2.0 (v2.2.7)^91^ was used to filter samples with contamination above 5%, using the contigs’ coverage and the full marker gene set to obtain high-quality genome bins. Quast (v5.2.0)^92^ was performed to examine the metrics of assemblies, such as N50, L50, total number of contigs, total length of the assembly (bp), and GC content (**Supplementary Table 1**). The clean reads from each sample were mapped to its high-quality assemblies to estimate the coverage of contigs in assembled genomes using BBMap, and the mean coverage for each genome was calculated using samtools (https://github.com/samtools/samtools). The taxonomy of assembled genomes was classified by gtdbtk (v2.1.1)^93^ with the database (release207_v2) in the “classify_wf” mode. **Supplementary Table 1** lists the taxonomy for each of the sequenced strains, including the taxonomies from both the Genome Taxonomy Database (GTDB) and NCBI Taxonomy Database.

Based on statistical metrics of the assemblies, a single representative genome for each species identified by gtdbtk (v2.1.1)^93^ was selected from the 1134 high quality isolated genomes; representative genomes were selected based upon assembly quality. dRep (v2.2.3)^94^ and fastANI (v1.32)^95^ were used to compare and calculate the average nucleotide identity (ANI) of the representative genomes of the 196 species. 95% ANI and 96% ANI cut-offs were used to cluster genomes into different groups (-pa 0.95 -sa 0.96 --S_algorithm fastANI). Phylogenomic analysis of the representative genomes of bladder strains were conducted using anvi’o v7^96^; assemblies were made into anvi’o contig databases (anvi-gen-contigs-database) and populated with hmms (anvi-run-hmms). The multiple sequence alignment of the concatenated amino acid sequences for 71 universal single-copy marker genes was extracted via the anvi’o command anvi-get-sequences-for-hmm-hits --get-aa-sequences --hmm-source Bacteria_71 --return_best-hit --concatenate. Phylogenetic trees were constructed using FastTree v2^97^ with default settings (JTT+CAT model) and visualized in the iTOL v6 web browser^98^. Genomes were single annotated using the Database of Clusters of Orthologous Genes (COGs 2020)^99^ and KEGG (Kyoto Encyclopedia of Genes and Genomes)^100^ in anvi’o v7^96^. Representative vaginal and gut genomes (**Supplementary Table 3**) also were selected and annotated as described above. The pan-genomes of bladder, vaginal and gut representative genomes were generated using the parameter (anvi-pan-genome) with frags (-- minbit 0.6 and --mcl-inflation 2). Using these pan-genomes, enrichment analysis for KEGG and COG functions was then conducted in anvi’o v7 with the parameter (anvi-compute-functional- enrichment) ^101^.

### Microbial 16S rRNA gene amplicon sequence analysis

16S rRNA gene sequence analysis of previously obtained 392 female bladder urine samples^3, 5, 35–37^ was conducted, as previously described^41, 42^. Cutadapt (cutadapt.readthedocs.io) was used to quality trim the raw reads derived from the 16S rRNA V4 region by removing adaptors from both ends (-a ^GTGCCAGCMGCCGCGGTAA…ATTAGAWACCCBDGTAGTCC -A ^GGACTACHVGGGTWTCTAAT…TTACCGCGGCKGCTGGCAC) and discarding processed reads that were shorter than 200 bp or longer than 240 bp (-m 200 -M 240 --discard-untrimmed). To generate amplicon sequence variant (ASV) tables, DADA2^102^ was applied to process the trimmed reads, including quality control, dereplication, and chimera removal. 392 bladder urine samples with >1000 ASV counts were kept for downstream analysis. We then used BLCA v2.2^103^ with the NCBI 16S Microbial Database to obtain taxonomic identities at the genus level.

### Phylogenetic grouping and virulence profiling of *E. coli* strains

We assembled and annotated publicly available gut *E. coli* isolates^40^ following the protocol listed above for the bladder isolates. *E. coli* strains of bladder and gut with high coverage (>20×) were used for downstream pan-genome and phylogenetic analyses. The phylogroups of *E. coli* strains were determined using ClermonTyping^104^. Anvi’o v7 was applied for the pan-genome analysis of *E. coli* strains using the parameter (anvi-pan-genome) with frags (--minbit 0.6 and -- mcl-inflation 10). Phylogenetic analysis for bladder *E. coli* strains was conducted by extracting amino acid sequences of 1084 single-copy core genes; amino acid sequences were concatenated and aligned (--concatenate-gene-clusters) in anvi’o v7. The phylogenetic tree was constructed and visualized as mentioned above. The genomes were screened for virulence factors and antibiotic resistance genes via abricate V.1.0.1 (https://github.com/tseemann/abricate) using the following databases: Virulence Factor Database^105^ (VFDB, updated 2021-Mar-27) and Comprehensive Antibiotic Resistance Database^106^ (CARD, updated 2021-Mar-27). Results were parsed via Python.

### Statistical analyses

All the statistical analyses were conducted using RStudio v3.6. The “ggplot2” R package was used for box plots, and “pheatmap” was used for heatmaps. Discriminant Analysis of Principal Components (DAPC) was conducted using “adegenet” package using ‘dapc’ function. Principal Coordinates Analysis (PCoA, function “capscale” with no constraints applied) was conducted using “vegan” package and the vegdist function with method ‘bray’ for bray-cutis dissimilarity analysis. Different groups in the ordination plot were tested using a PERMANOVA test (function “adonis”) and were clustered on the plot using function “ordiellipse” with statistics (kind = “sd”, conf = 0.95).

### Data availability

Raw data of whole genome sequence for isolates and assemblies of isolates are available in the Sequence Read Archive under BioProject PRJNA970254. The BioProject will be released to the public upon the publication of the paper. Refer to Alan Wolfe or Jingjie Du for the availability of the assemblies of bladder isolates and gut *E. coli* isolates. Supplemental Table 1 includes taxonomy, assembly statistics, study information (participant number, study code, sample type) and participant information (age, gender, race BMI, symptom status).

## Supporting information

https://cornell.box.com/s/35xc0k6zw6y9ozv89caxmkdgmpp6bsni

## Acknowledgements

This work was funded by multiple sources, including R01DK104718 from the National Institute of Health, a contract from Pathnostics, and a gift from an anonymous donor. We wish to acknowledge the study participants who consented to donate urine and the clinicians who recruited those participants and collected their urine. We especially wish to acknowledge the clinical members of the Loyola Urinary Education and Research Collaborative (LUEREC), who obtained the vast majority of urines associated with this study. We thank all the members of the Wolfe laboratory who processed the urine samples to identify isolates and Genevieve Baddoo for isolation of a few of the strains included in this study. Finally, we wish to thank Paul Schreckenberger (deceased) and Amanda Harrington, former and current directors of the Loyola University Clinical Microbiology Laboratory for their willingness to share their MALDI-TOF instruments.

---

**Supplementary Figure 1:**
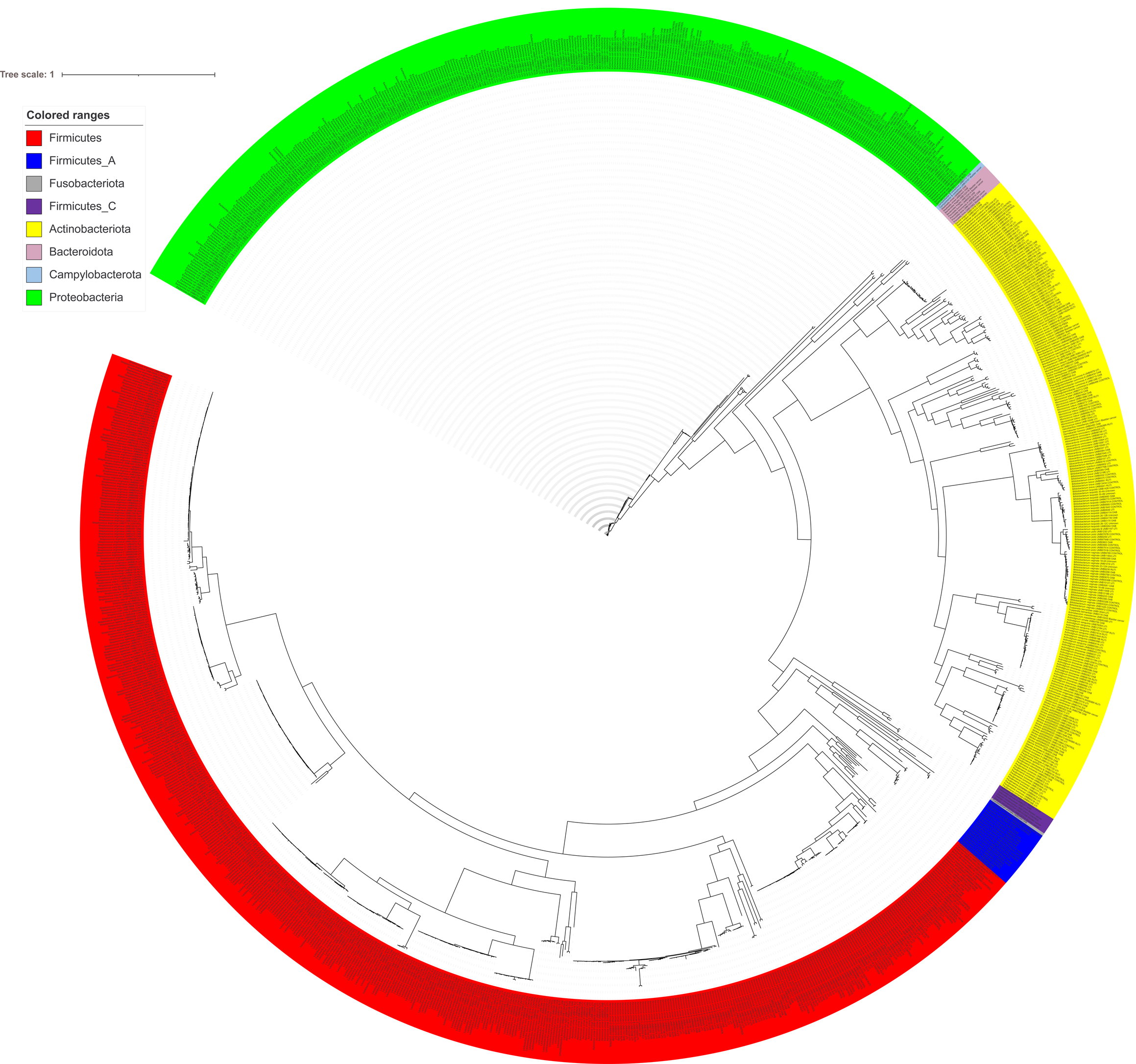
Phylogenomic tree of 1134 bladder isolate genomes. 71 core universal bacterial genes were used to reconstruct the phylogenetic tree. Bacterial species are colored by GTDB-identified phylum. Disease status of the host for each isolate is indicated in the isolate’s name.

**Supplementary Figure 2:**
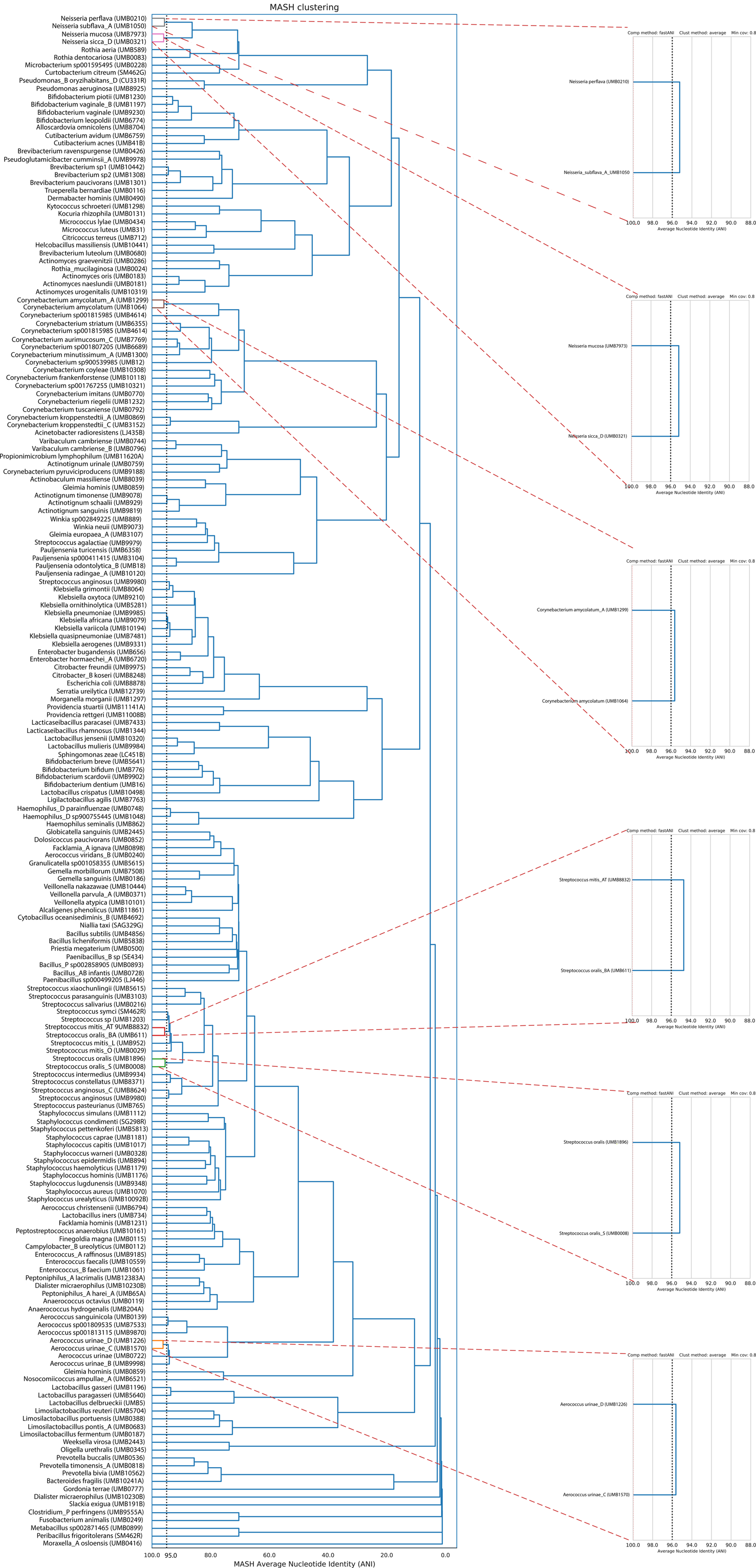
Whole-genome ANI comparison of the 196 different GTDB-identified bacterial species in the bladder isolate genomes.

**Supplementary Figure 3:**
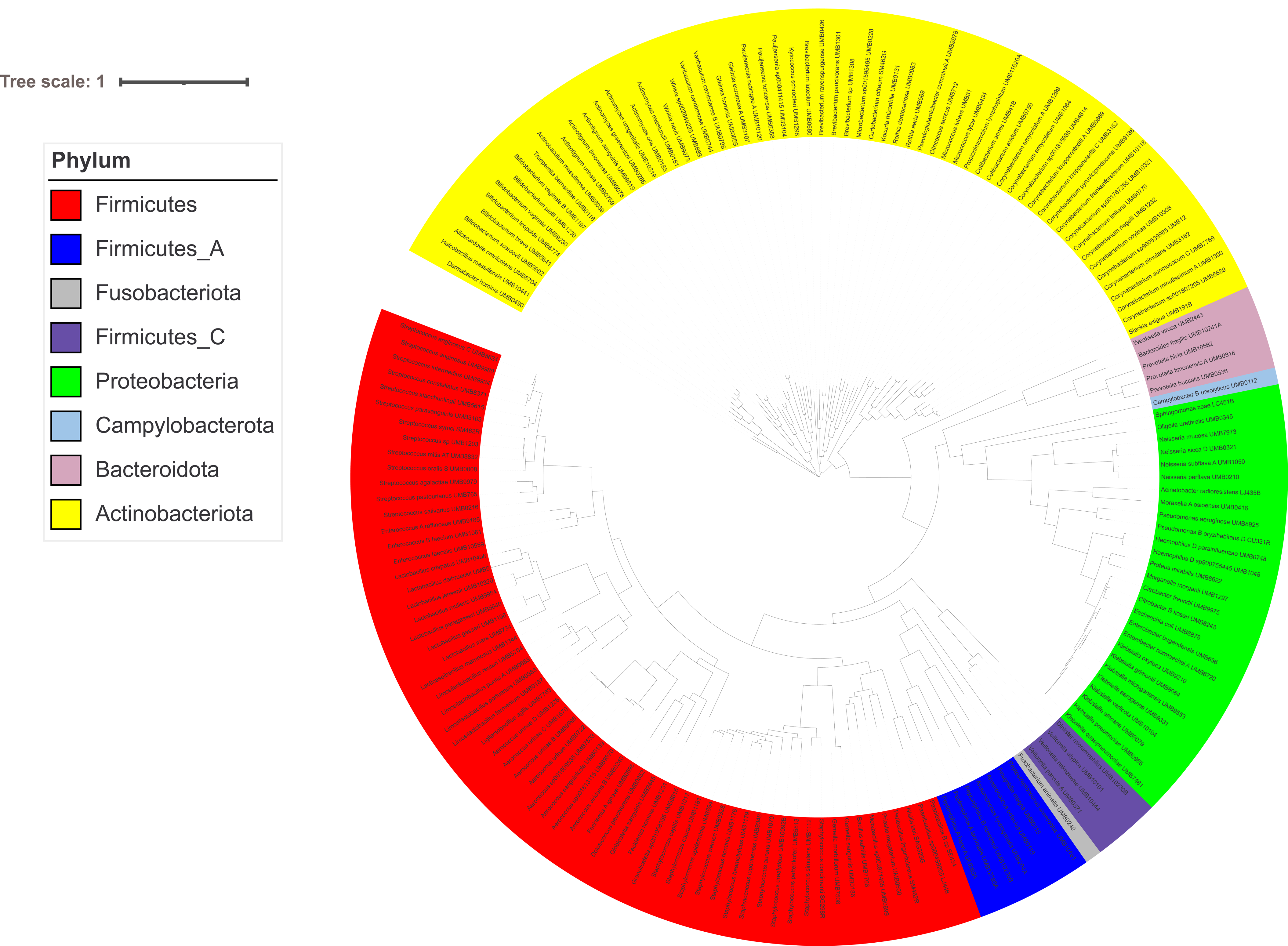
Phylogenomic tree of 169 bacterial species (assigned by GTDB) represented in the bladder genome collection from the symptomatic group. A single representative genome per species was selected out of the 837 high quality isolated genomes (>90% completeness, < 5% contamination) isolated from the symptomatic group to reconstruct the multiple sequence alignment of 71 core bacterial genes. Bacterial species are colored by phylum.

**Supplementary Figure 4:**
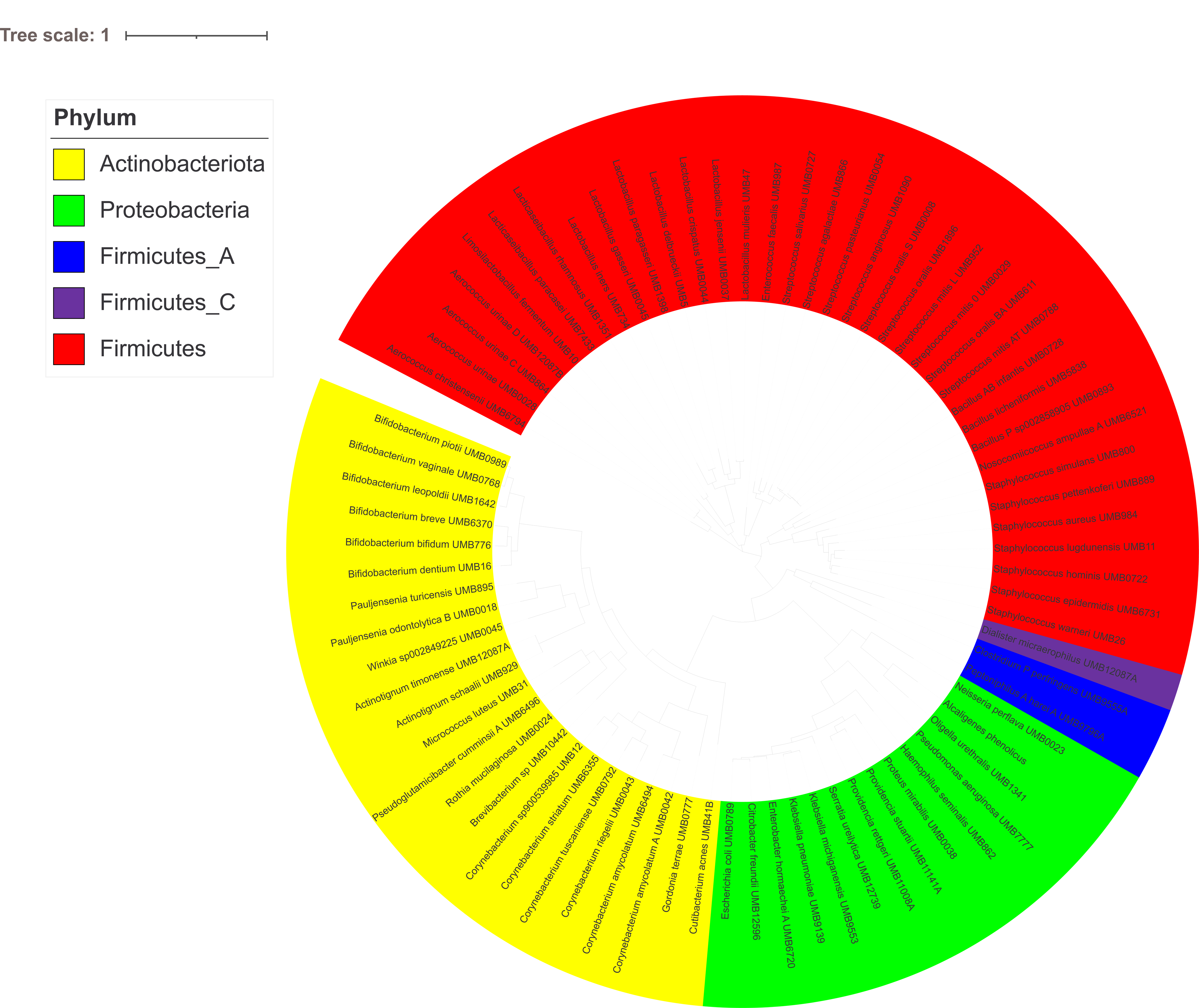
Phylogenomic tree of 76 bacterial species (assigned by GTDB) represented in the bladder genome collection from the asymptomatic group. A single representative genome per species was selected out of the 239 high quality isolated genomes (>90% completeness, < 5% contamination) isolated from healthy group to reconstruct the multiple sequence alignment of 71 core bacterial genes. Bacterial species are colored by phylum.

**Supplementary Figure 5:**
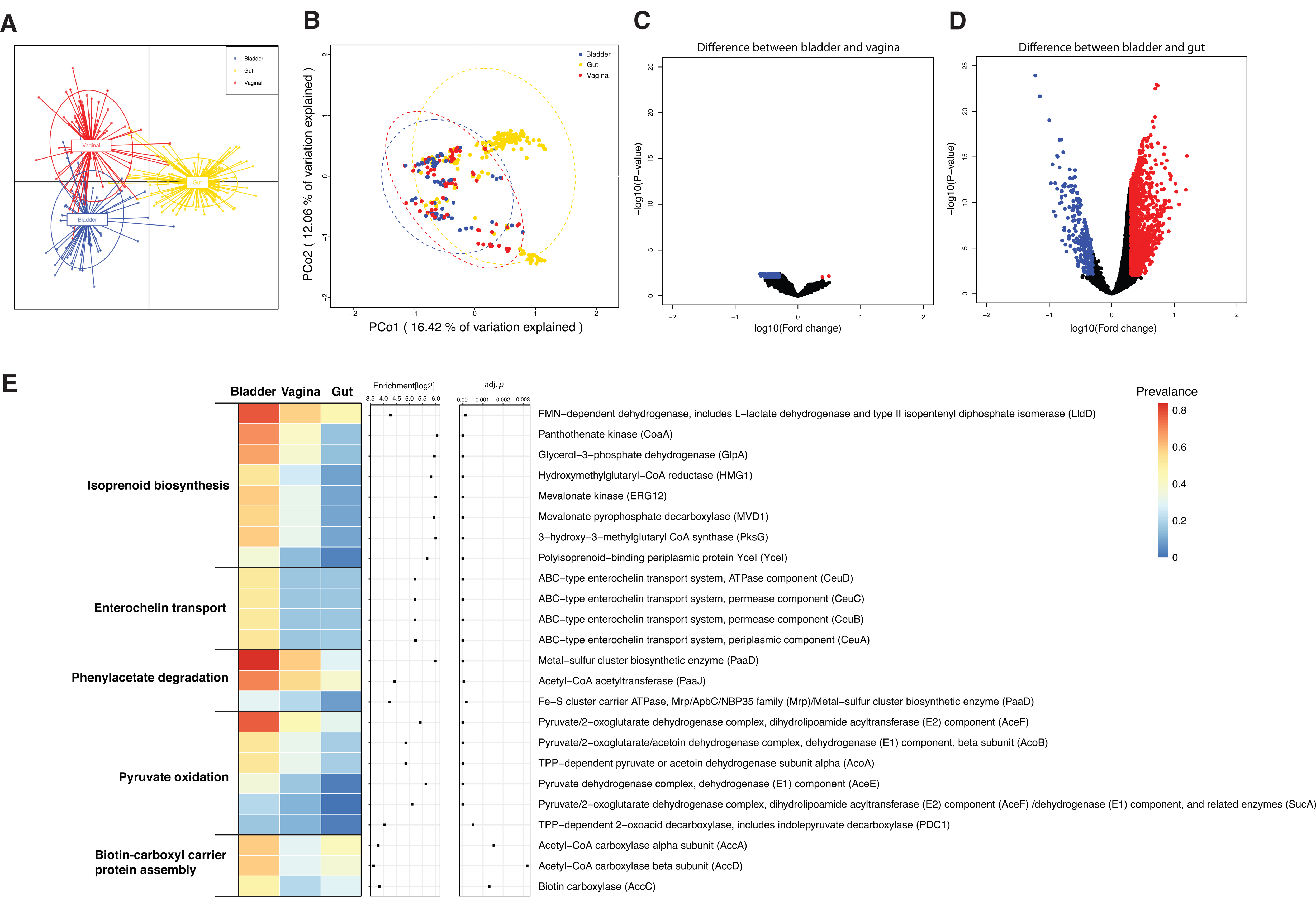
COG function comparisons of bladder isolate genomes with vaginal and gut isolate genomes. A-B. DAPC and PCoA of COG functional diversity of representative bacterial species isolated from the asymptomatic individuals of different niches. D. Volcano plots of *t* tests corrected by the Benjamini and Hochberg method for changes of COG functions between bladder and gut/vagina. An FDR cutoff of 0.01 and [logFC] > 0.3 were used. Data points highlighted in red indicate COG functions that were significantly enriched in gut or vagina, while data points highlighted in blue indicate COG functions that were significantly enriched in the bladder. F. Selected COG functions that were enriched in the bladder as compared with gut and/or vagina. An FDR cutoff of 0.01 and more than 10% prevalence in all bladder genomes were used.

**Supplementary Figure 6:**
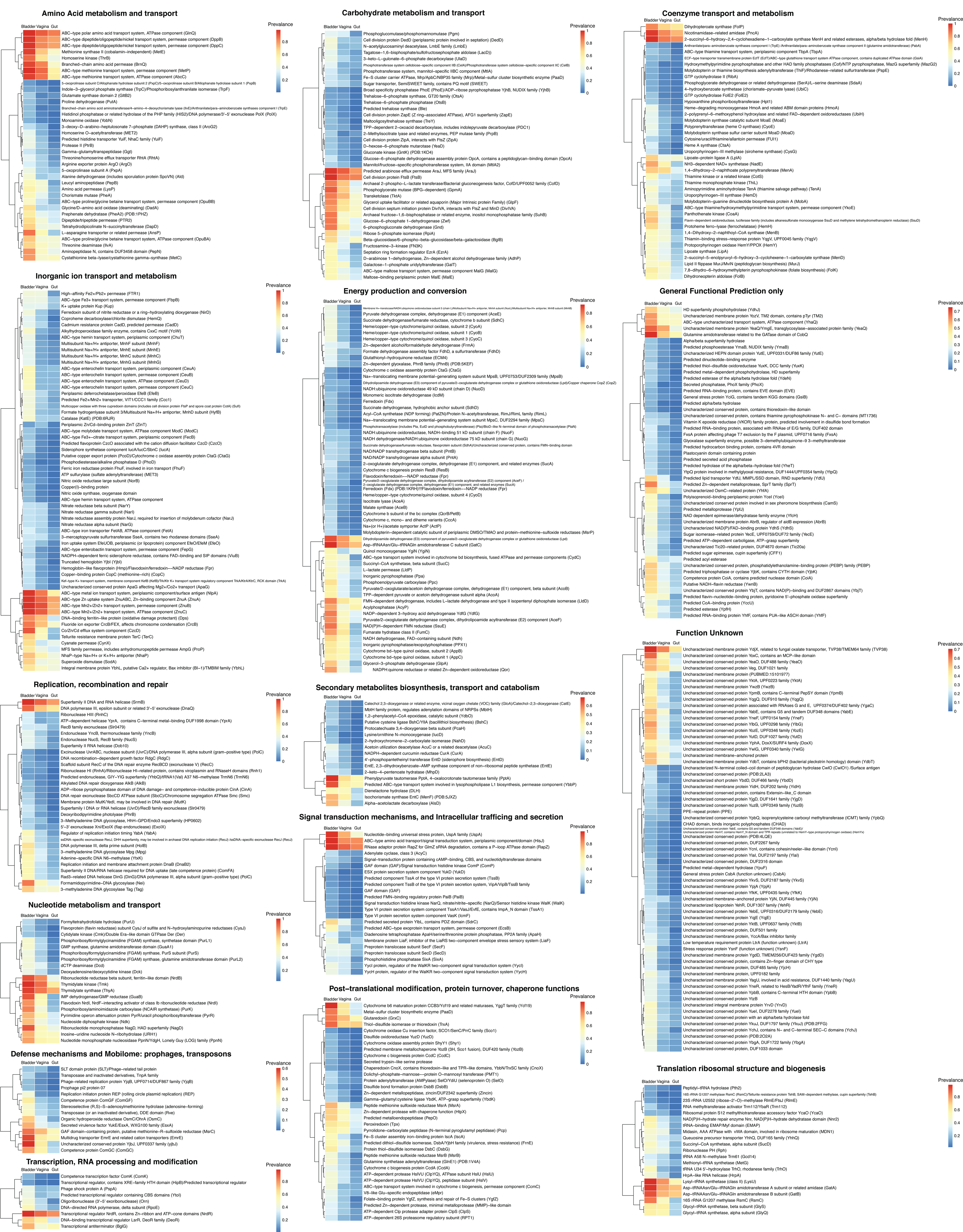
COG functions enriched in the bladder based on COG category. COG functions with an FDR cutoff of 0.01 and [logFC] > 0.3 were displayed here.

**Supplementary Figure 7:**
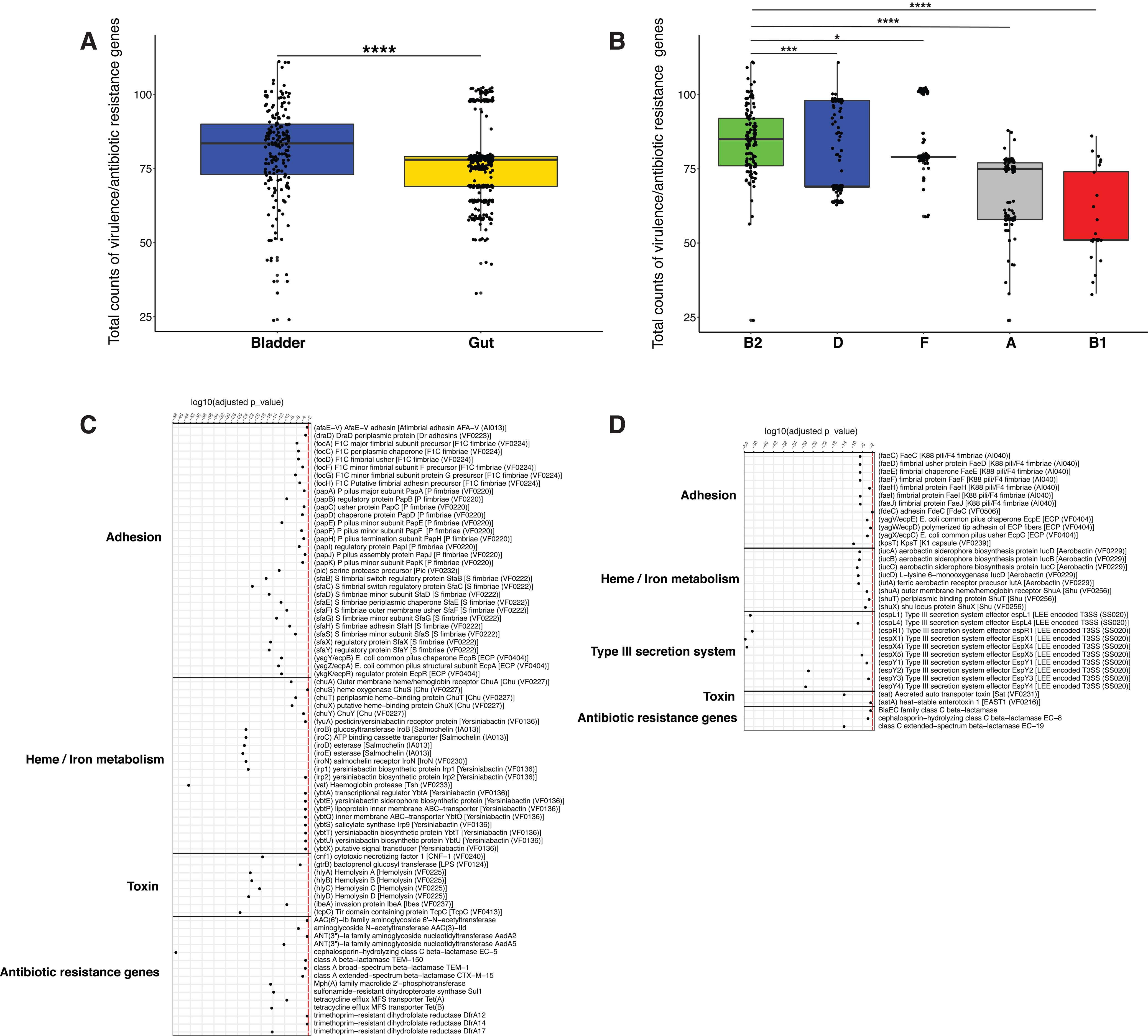
Comparison of virulence and antibiotic resistance genes between *E. coli* genomes isolated from bladder and gut. A. Boxplots analysis showing the comparison of counts of virulence and antibiotic resistance genes in bladder and gut *E. coli* genomes. Each data point represents the counts of virulence and antibiotic resistance genes in each *E. coli* genome. B. Boxplots analysis showing the comparison of counts of virulence and antibiotic resistance genes in different *E. coli* phylogroups. Phylogroup G was not shown here due to low sample size. *, **, ***, **** indicated FDR<0.05, 0.01, 0.001 and 0.0001 respectively. C. Virulence and antibiotic resistance genes that were enriched in bladder *E. coli* genomes. D. Virulence and antibiotic resistance genes that were enriched in gut *E. coli* genomes. Virulence and antibiotic resistance genes with an FDR of <0.01 were displayed here.

## Supplementary Tables

**Supplementary Table 1:** Metadata for bladder-specific isolated genomes and hosts information.

**Supplementary Table 2:** Relative abundances of different genera in 16S sequencing samples.

**Supplementary Table 3:** List of GTDB classified species isolated from gut, vaginal, and bladder samples. A single representative genome was selected for each species from the gut, vaginal, and bladder strains.

**Supplementary Table 4:** KEGG functions of bladder, gut, and vaginal representative genomes.

**Supplementary Table 5:** COG20 functions of bladder, gut, and vaginal representative genomes.

**Supplementary Table 6:** Statistical analyses of KO modules in bladder, vaginal, and gut representative genomes. KO module enrichment table was generated using anvi’o v7.

**Supplementary Table 7:** Statistical analyses of KOfam hits in bladder, vaginal, and gut. Representative genomes. KOfam enrichment table was generated using anvi’o v7.

**Supplementary Table 8:** Statistical analyses of COG20 functions in bladder, vaginal, and gut representative genomes. COG20 function enrichment table was generated using anvi’o v7.

**Supplementary Table 9:** Phylogroup distribution of bladder and gut *E. coli* strains.

**Supplementary Table 10:** KOfam profiles of bladder *E. coli* strains.

**Supplementary Table 11:** KOfam profiles of gut *E. coli* strains.

**Supplementary Table 12:** Differentially abundant KOfam profiles between bladder and gut *E. coli* isolates by Wilcoxon rank sum test.

**Supplementary Table 13:** Virulence factor genes identified in bladder *E. coli* isolate assemblies. Number listed is number of copies of the gene.

**Supplementary Table 14:** Antibiotic resistance genes identified in bladder *E. coli* isolate assemblies. Number listed is number of copies of the gene.

**Supplementary Table 15:** Virulence factor genes identified in gut *E. coli* isolate assemblies. Number listed is number of copies of the gene.

**Supplementary Table 16:** Antibiotic resistance genes identified in gut *E. coli* isolate assemblies. Number listed is number of copies of the gene.

**Supplementary Table 17:** Differentially abundant virulence and antibiotic resistance genes between bladder and gut *E. coli* isolates by Wilcoxon rank sum test.

